# On the Independence of Individual Elements of Visual, Auditory and Tactile Sensor Structures in the Functioning of a Simple Sensorimotor Reaction

**DOI:** 10.64898/2026.01.27.701918

**Authors:** Alexey A. Kulakov

## Abstract

**Relevance:** For performance in a dynamically changing environment, not only the speed of a sensorimotor reaction is important, but also the speed of its recovery (relaxation) after a previous response.

**Goal:** To investigate whether the neural pathways from various receptors or their groups are functionally independent in the process of recovery after excitation transmission.

**Methods:** In over 20 subjects, the latent period of a simple sensorimotor reaction (SSMR) to visual, auditory, and tactile stimuli was recorded with varying interstimulus intervals. The relaxation parameters of the variable component of the reaction time were extracted by approximating the data with a multiexponential model. The key paradigm involved alternating stimulation of different sensors: spectrally different (red/blue light), totally different (sounds of different frequencies), or spatially separated (different areas of the retina/skin).

**Results:** It was shown that alternating stimulation, compared to isolated stimulation of a single type/location, leads to a significant reduction in the time constants of SSMR relaxation. The effect was revealed for all studied modalities.

**Conclusion:** The obtained data indicate the functional independence of neural channels processing information from different receptors or their groups during the recovery phase after excitation transmission, up to the level of the motor center. This suggests a higher degree of specificity in the organization of sensorimotor responding than might be assumed based on data about the diffuse nature of cortical activation recorded by EEG and fMRI methods.

## Introduction

The presence of paired sensory systems is not only an important evolutionary acquisition of animals but also holds certain value for their study, as one of them can always serve as a reference system when studying the effects of various external factors. For this reason, the study of hearing and vision has achieved significant success.

The structure of these systems has shown that the directly sensing elements in these systems are quite numerous, each contributing to the creation of a visual or auditory image.

In the case of vision, these are the light-sensitive cones and rods. Rods have the greatest sensitivity in the green region of the spectrum, while the three types of cones have peak sensitivities in the blue, green, and red regions of the spectrum. Their sensitivity ranges overlap, providing continuous sensitivity from 400 nm to 700 nm [1]. The total number of cones is 6-7 million [2].

In the case of hearing, these are the hair cells of the organ of Corti, numbering almost 20,000 cells, each tuned to a specific oscillation frequency. Each of these cells transmits excitation to its nerve cell, which ultimately reaches the visual and auditory fields of the cerebral cortex [3].

In the case of touch, this is the most extensive receptive field – the skin. The skin contains 4 types of tactile receptors responding to touch, pressure, slip, and vibration [4]. Information from tactile receptors is then transmitted from the skin to the upper spinal cord and further to the thalamus, primary and secondary somatosensory areas, which are the endpoint of a hierarchical organization with various overlapping networks involved in performing different functions [5].

Thus, in all cases, information about the occurrence of excitation in a receptive field ultimately reaches the corresponding sections of the brain. This raises the question: if we excite only a part of the sensory cells, will the entire sensory field in the cortex be excited? According to EEG data, excitation spreads to significant parts of the cortex [6-13]. fMRI observations show that excitation has a more localized character but also occupies sufficiently large areas [14-23].

An affirmative answer would mean that, regardless of the initial excitation of a few sensory cells, excitation of the majority of neurons occurs in the receptive field (auditory, visual, etc.) with guaranteed transmission to motor areas.

To test this assertion, we utilized one feature of the simple sensorimotor reaction (SSMR) that has been undeservedly overlooked by researchers. This is the dependence of the SSMR latent period on the waiting time after the previous response. It appears that with an increase in waiting time, the latent time gradually decreases to some stationary level [24-26]. It has been shown that this decline is satisfactorily described as exponential [27] and even multiexponential [26], i.e., as a process developing over time. Consequently, this process can be influenced in one way or another.

In this context, we have shown [28] that with alternate excitation of paired sensors (visual or auditory), a decrease in the relaxation constant of the simple sensorimotor reaction after responding to a signal is observed. This apparent acceleration was explained by the fact that during the response, when one eye (ear) is excited, the reflex arc of the other eye (ear) restores its ability to quickly transmit excitation to the motor area. This assumption contradicts the aforementioned view of diffuse excitation of the majority of neurons in the corresponding receptive field in the cerebral cortex.

This article is, as it were, a continuation of our publication “Is the Simple Sensorimotor Reaction Simple?”. Here we present data on the relaxation rate in the case of alternating excitation of parts of visual and auditory receptors and put forward the following hypothesis: “the components of the simple sensorimotor reaction reflex arc, originating from individual types of receptors or individual areas of the receptor field, are functionally independent in terms of temporal recovery (relaxation) after excitation conduction.

## Methods

The studies involved 3rd-4th year students, as well as staff of the KNITU-KAI department (more than 20 subjects). Measurements were carried out using a computer program, as well as on a hardware-computing complex including an Arduino Nano microcontroller and a computer.

The participant was asked to respond to the appearance of a sound or light signal by pressing a button or tapping on the table with a pen. The course of the study was cyclical: after the previous response, a waiting interval was set by the program, after which a signal, acoustic or light, was presented. The participant responded, and the cycle repeated. The signal presentation program was a Greco-Latin square, so that the sum of the presentation order was approximately equal vertically and horizontally. The number of presentations was 50 or 2x25. Thus, the factor of the waiting interval sequence was minimized, and the subject could not predict the length of the next waiting interval for the signal. The Greco-Latin square was created anew each time. The light signal was presented as a spot on the computer screen, blue or red, or as the glow of an LED, blue or red, as well as white, and continued until the participant responded. Blue or red colors were chosen to excite the corresponding cones in the subjects’ retinas, as their sensitivity spectra practically do not overlap [1]. The luminance of the red spot on the display screen was 75 cd/m^2^, blue - 86 cd/m^2^. The luminance of white LED light was 500 cd/m^2^, red - 100 cd/m^2^, blue - 280 cd/m^2^. The sound signal was synthesized on the Arduino Nano microcontroller as a square wave with frequencies of 932, 880, 988, 1047 Hz, which approximately corresponded to A, A#, B of the second octave, and C of the third octave (i.e., the “BACH” theme). Sound intensity was on average 75 dB. In studies with Arduino, PKM-1B buttons with a magneto-reed switch were used. In studies of the tactile-motor reaction, the Arduino-computer system was also chosen. Vibration motors from smartphones (both as motors with an eccentric and as “pills”) were used to elicit the reaction, attached to the skin with adhesive tape. SSMR calculations were performed as in [26,28,29]. We used a modified version of Kukinov’s technique [30]. The sequence of calculations is illustrated in Fig. 1.

**Figure 1.**
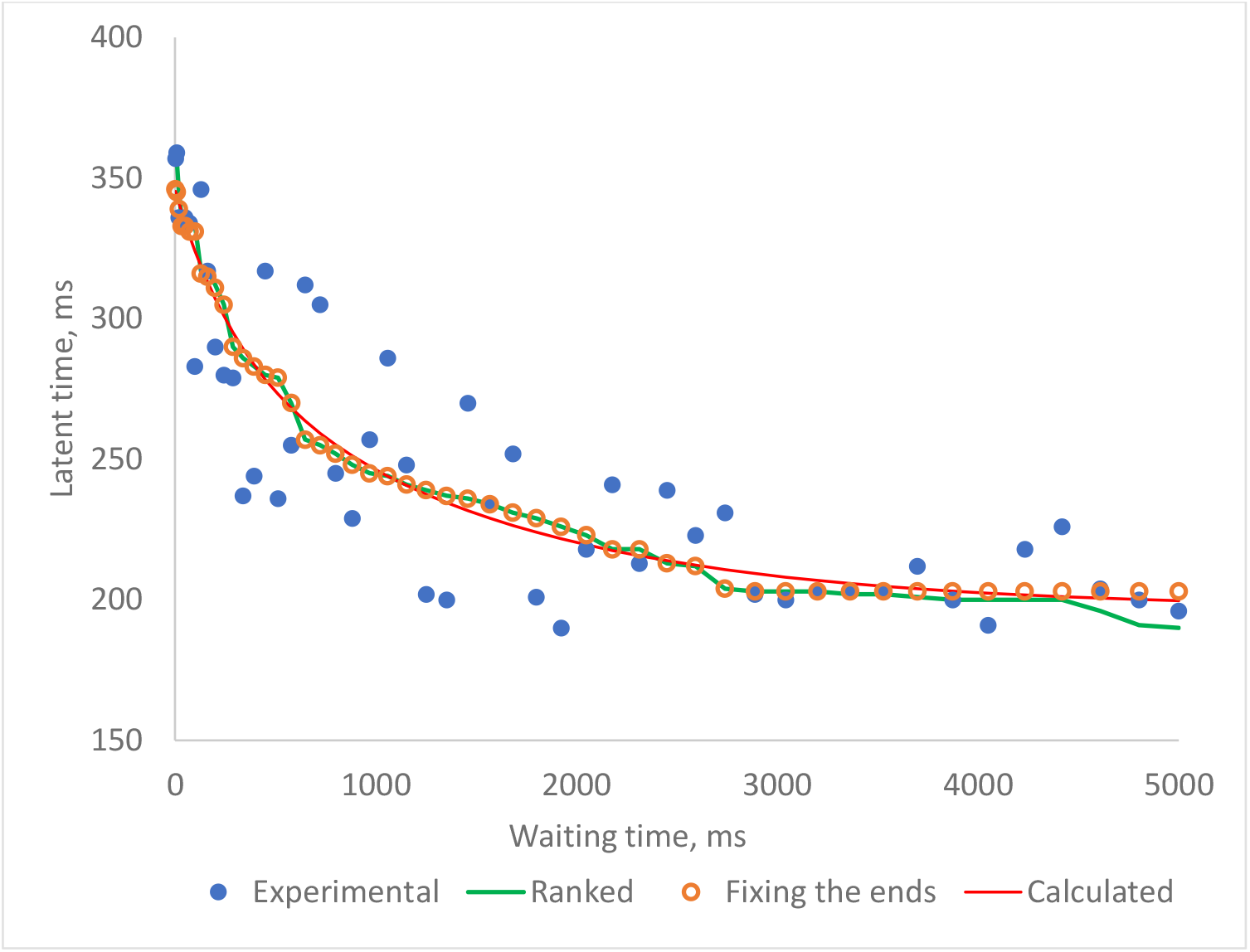
Stages of data modification for SSMR calculation: a) Blue circles - experimental data; b) after ranking the experimental data; c) orange circles - fixed ends; d) calculated dependence.

Briefly, it happened as follows. 1. Initially, experimental data were taken. 2. These latent time data were ranked in descending order. 3. At the end of the dependence, at approximately 80% of the waiting time, where the decline of the dependence began after some delay, the data were fixed at one level. This difference was symmetrically subtracted from the first points at the beginning of the dependence. This constituted the modification of Kukinov’s method [30], who believed that after ranking, the entire curve should be rotated slightly counterclockwise, removing the contribution of errors. Thus, the error was extracted, which as a result of ranking was also arranged in a variational series, with the most deviating from the main dependence mainly at the edges. 4. The obtained data were entered into Excel, where an approximation of these data in the form of a multiexponential decay of 3 exponents was performed according to the Formula:

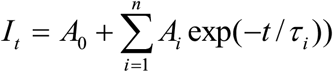

where A0 is the constant value of latent time, Ai and τi are the amplitude and relaxation time of the i-th component, respectively, t is the time interval from the previous reaction to irritation, I(t) is the experimental value of the SSMR latent time, and n=3. The value n=3 best described the dependence (in reality, we often discarded the fastest component later due to its small contribution to the dependence). During the calculation, the values of amplitudes and relaxation times were selected using the criterion R^2^ = Σ(I_exp_ - I_calc_)^2^, where I_exp_ and _alc_ are the values of the ranked and calculated latent times, respectively, using the conjugate gradient method for the entire system of equations.

## Results

A feature of the simple sensorimotor reaction is its variability. First, in the range of 0-5 sec of stimulus waiting, the reaction latent time is initially quite long – up to 1-2 sec, and then decreases quite quickly to a fairly constant value [24-26], with the decline in latent time well described by a multicomponent exponential [26]. One of the explanations for this phenomenon is a) the occurrence of inhibition b) depletion of resources for new excitation [26,31]. Second, over a sufficiently long time, the components of the reaction latent time change quite significantly [29]. Based on these assumptions, we showed that alternating excitation of paired sensors leads to an acceleration of the relaxation of the decline in the variable part of the latent time [28]. This happened, as it seemed to us, because when one sensor is excited, for example, the right one, the left sensor at that time restored its ability to respond. In this regard, the idea arose to check whether individual areas of the retina could behave similarly. We investigated the effects of the color of spots (red, blue, and alternating blue-red) on the display screen on the magnitude of the latent time of the simple sensorimotor reaction. Table 1 presents the results of this experiment.

**Table 1.** The effect of a light stimulus of different colors on the relaxation parameters of a simple sensorimotor reaction.

| Subject | color of stimulus | Relaxation parameters, ms |  |  |  |  |
| --- | --- | --- | --- | --- | --- | --- |
|  |  | A<sub>1</sub> | T<sub>1</sub> | A<sub>2</sub> | T<sub>2</sub> | A<sub>0</sub> |
| A1 | Red | 114.6 | 34.6 | 74.4 | 1358.4 | 249.5 |
|  | Red-Blue | 77.0 | 16.8 | 82.1 | 881.5 | 230.9 |
| A2 | Red | 123.4 | 21.9 | 120.6 | 1308.8 | 238.4 |
|  | Red-Blue | 49.2 | 101.9 | 52.5 | 2030.6 | 207.5 |
| A3 | Red | 238.4 | 57.1 | 104.6 | 1010.1 | 244.2 |
|  | Red-Blue | 59.2 | 132.4 | 64.6 | 1285.8 | 256.7 |
| I1 | Red | 84.0 | 200.3 | 95.6 | 1763.9 | 252.5 |
|  | Red-Blue | 95.5 | 1.1 | 46.4 | 949.4 | 236.0 |
| I2 | Red | 63.1 | 5.6 | 144.2 | 1021.5 | 259.1 |
|  | Red-Blue | 26.8 | 6.4 | 73.4 | 1205.0 | 273.0 |
| K1 | Red | 107.9 | 28.6 | 144.3 | 654.2 | 241.0 |
|  | Red-Blue | 96.9 | 32.0 | 75.3 | 1549.3 | 229.3 |
| K2 | Red | 183.3 | 5.4 | 62.1 | 941.9 | 242.4 |
|  | Red-Blue | 45.1 | 130.9 | 65.0 | 1318.4 | 252.2 |
| K3 | Red | 113.3 | 1.1 | 136.4 | 644.2 | 214.4 |
|  | Red-Blue | 4.6 | 3.5 | 66.8 | 853.1 | 215.8 |
| K4 | Red | 90.0 | 36.8 | 120.9 | 550.7 | 230.4 |
|  | Red-Blue | 57.0 | 75.9 | 61.8 | 1215.0 | 228.3 |

The experiment was conducted using a computer. A square spot of red or alternating red and blue appeared on the dark background screen, approximately 0.8 cm in size.. The subject reacted to the appearance of a spot by pressing the mouse button, regardless of the color of the spot. The subjects were 18-19 years old

For one subject, we conducted a more thorough study. The results are presented in Table 2.

**Table 2.** Changes in the parameters of a simple sensorimotor reaction when stimulated by a spot with one color, and alternately with two.

| Relaxation parameters,<br>ms | Reaction conditions: the color of the spot |  |  |
| --- | --- | --- | --- |
|  | Blue | Red | Red-Blue |
| A<sub>1</sub> | 93.9 ± 17.2 | 84.6 ± 11.8 | 80.6 ± 15.1 |
| T<sub>1</sub> | 65.5 ± 22.2 | 57.9 ± 13.3 | 24.7 ± 16.0 |
| A<sub>2</sub> | 64.6 ± 7.2 | 72.8 ± 13.5 | 89.9 ± 11.1 |
| T<sub>2</sub> | 1305.7 ± 119.3 | 1261.9 ± 228.8 | 528.8 ± 108.0 |
| A<sub>0</sub> | 256.9 ± 2.7 | 250.2 ± 2.7 | 255.8 ± 5.3 |

From Table 1, it can be seen that the result was ambiguous: the relaxation time constant for blue-red excitation was both less and greater than that for a constant color spot. It did not matter whether the background was light or dark. Analyzing the results, we concluded that: 1. The spot sizes were small, and due to floating gaze fixation, an inconstant part of the retina was illuminated. 2. The subject saw the spots with both eyes, which smoothed out the differences. 3. The intensity of the spots was not sufficiently different from the background, i.e., the light did not excite all cones of the illuminated part of the retina, which reduced the impact. 4. We have shown that during measurements, the relaxation parameters of the variable part of the latent time can change quite significantly [29].

In this regard, we switched to using LEDs and a hardware-measuring complex (Arduino Nano + computer). Furthermore, we conducted only 1-2 measurement cycles per test session to prevent a strong shift in the psychological state of the subject and their fatigue. Finally, we used excitation of only one eye, with the subject focusing their gaze on an object (a non-luminous LED) located 3-4 angular degrees from the luminous LED. We assumed that this would create conditions for exciting the same area of the retina. The results of these measurements are shown in Table 3.

**Table 3.** Comparison of the excitation of a simple sensorimotor reaction by alternating blue-red light and monochrome LED lighting.

| Subjects | Relaxation parameters, ms | Reaction conditions: the color of the leds |  |  |
| --- | --- | --- | --- | --- |
|  |  | Blue | Red | Red-Blue |
| AK4, 74,m | A<sub>1</sub> | 47.0 ± 1.4 | 73.1 ± 2.0 | 80.6 ± 0.6 |
|  | T<sub>1</sub> | 303.4 ± 16.7 | 460.2 ± 15.6 | 84.4 ± 0.5 |
|  | A<sub>2</sub> | 33.2 ± 1.1 | 14.0 ± 2.3 | 34.1 ± 2.0 |
|  | T<sub>2</sub> | 972.4 ± 43.9 | 1647.0 ± 303.7 | 820.4 ± 55.3 |
|  | A<sub>0</sub> | 244.6 ± 1.0 | 243.0 ± 1.8 | 239.7 ± 1.2 |
| L,45,m1 | A<sub>1</sub> | 97.1 ± 4.2 | 108.4 ± 29.14 | 133.3 ± 24.1 |
|  | T<sub>1</sub> | 290.5 ± 135.9 | 54.8 ± 25.5 | 99.7 ± 70.6 |
|  | A<sub>2</sub> | 26.0 6. ± 3 | 55.4 ± 3.9 | 133.2 ± 31.4 |
|  | T<sub>2</sub> | 1277.7 ± 186.0 | 1039.9 ± 53.2 | 506.6 ± 12.9 |
|  | A<sub>0</sub> | 196.7 ± 0.9 | 197.9 ± 4.7 | 188.4 ± 4.6 |

The experiment was performed similar to Tables 1,2, but with LEDs as stimulating signals. For the AK4 subject, there was a significant difference in alternating red-blue excitation in T2 ( $p<0.03$ ) as compared with red excitation, and for T1 – with both red and blue. For subject L1, there was a significant difference for T1 (both for red and blue light), T2 and Tau ( $p<0.01$ ) between alternating red-blue excitation and monochrome red and blue.

From the Table 3, it can be seen that when the eye is excited alternately with blue and red light, the relaxation constants, its main, slow component, are significantly less than when excited only with red or blue light. The same applies to the faster component. At the same time, the value of the constant component of the latent time decreases slightly.

A feature of excitation with LEDs is significantly greater brightness and, accordingly, contrast compared to experiments on computers. Therefore, we tested the experiment with excitation by bright and weak light. The data are presented in Table 4.

**Table 4.** The effect of different excitation colors on the parameters of a simple sensorimotor reaction: bright and weak excitation.

| Stimulation light | Relaxation parameters, ms | Low brightness | Full brightness |
| --- | --- | --- | --- |
| Blue | A<sub>1</sub> | 58.9 ± 5.1 | 67.5 ± 7.2 |
|  | T<sub>1</sub> | 133.6 ± 22.6 | 164.3 ± 63.9 |
|  | A<sub>2</sub> | 54.0 ± 3.9 | 70.2 ± 16.4 |
|  | T<sub>2</sub> | 1510.5 ± 99.7 | 1334 ± 125.6 |
|  | A<sub>0</sub> | 228.6 ± 2.1 | 200.5 ± 3.8 |
| Red | A<sub>1</sub> | 59.6 ± 5.6 | 69.1 ± 10.0 |
|  | T<sub>1</sub> | 122.1 ± 46.4 | 220. 7 ± 32.1 |
|  | A<sub>2</sub> | 64.9 ± 4.9 | 56.7 ± 7.1 |
|  | T<sub>2</sub> | 1286.6 ± 78.7 | 1869.4 ± 241.6 |
|  | A<sub>0</sub> | 228.2 ± 1.9 | 200.1 ± 4.0 |
| Blue-Red | A<sub>1</sub> | 102.3 ± 18.9 | 87.9 ± 16.4 |
|  | T<sub>1</sub> | 43.2 ± 5.4 | 63.3 ± 29.6 |
|  | A<sub>2</sub> | 58.8 ± 4.4 | 82.0 ± 14.9 |
|  | T<sub>2</sub> | 960.1 ± 70.9 | 672.3 ± 67.7 |
|  | A<sub>0</sub> | 227.1 ± 1.8 | 206.7 ± 3.3 |

In this experiment, subject AK4, male, 74 years old, reacted to light from LEDs at full brightness (bright light) and low brightness (approximately 1% of the brightness). Subject AK4 (74), (m). Significant differences in relaxation times for alternate stimulation and single-color stimulation are indicated in bold. There are significantly high values of A<sub>0</sub> for low brightness relative to high brightness, but they are not marked for clarity.

From the table, it can be seen that the effect of accelerating the relaxation constant persists even with weak light excitation, but to a lesser extent than with bright light. However, a noticeable decrease in the magnitude of the constant component of the SSMR is observed.

Another variant of the simple visuo-motor reaction is the spatial arrangement of light sources. It is one thing to excite a specific place on the retina, another to do it in several places alternately. Then the place that is not subjected to excitation seems to “rest,” “recover.” And then the SSMR relaxation constants will appear shorter. Table 5 illustrates this.

**Table 5.**
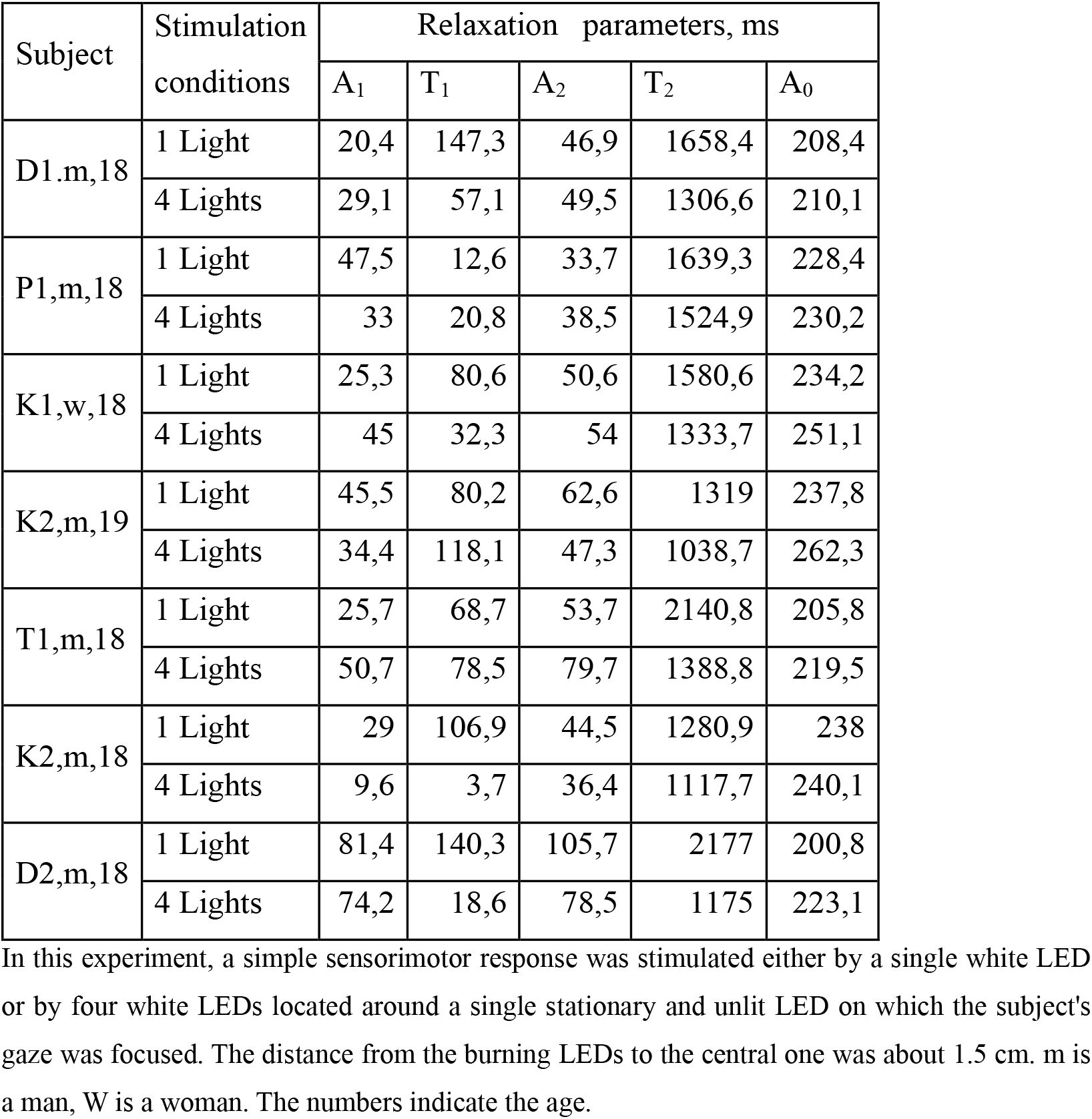
Changes in parameters of a simple sensorimotor response when stimulated by 1 or 4 LEDs.

The data in Table 5 represent the results of a single experiment and, therefore, the relaxation times T2 are not always noticeably shorter for the variant of excitation by alternating SSMR, the same applies to T1. Therefore, we undertook a more thorough study with a smaller number of subjects. The data are presented in Table 6.

**Table 6.** Effect of stimulation by one or alternately four light sources on the parameters of the simple sound-motor reaction (SSMR)

| Stimulation conditions | Relaxation parameters, ms | Subject AK4, m,73 | Subject V1,w,34 |
| --- | --- | --- | --- |
| 1 Light | A<sub>1</sub> | 53.9 ± 9.9 | 93.5 ± 7.6 |
|  | T<sub>1</sub> | 149 ± 49.8 | 283.0 ± 52.4 |
|  | A<sub>2</sub> | 71.8 ± 5.3 | 51.9 ± 9.6 |
|  | T<sub>2</sub> | 1588.5 ± 230.7 | 1562.9 ± 125.8 |
|  | A<sub>0</sub> | 236.2 ± 9.32 | 201.6 ± 1.8 |
| 4 Lights | A<sub>1</sub> | 55.2 ± 10.9 | 99.2 ± 17.6 |
|  | T<sub>1</sub> | 20.7 ± 6.7 | 62.8 ± 20.4 |
|  | A<sub>2</sub> | 128.9 ± 13.3 | 89.9 ± 10.5 |
|  | T<sub>2</sub> | 679.5 ± 68.0 | 793.3 ± 84.8 |
|  | A<sub>0</sub> | 215.9 ± 2.0 | 208.3 ± 2.4 |

The experiment was conducted as described in Table 6. However, the number of tests consisted of 5-6 series on different days. one episode per day. For a subject with AC4. the difference between the options Where the difference between the corresponding parameters was significant for this subject, the values of these parameters were indicated in bold

As we can see, with alternating excitation of different areas of the retina, the apparent relaxation rate increases. Next, we tried to test this effect for sound. Here we increased the sound intensity compared to [29] to 75 dB. We applied excitation with sounds of different pitch. A control sound of 932 Hz was used, and as an alternative, 4 cyclically changing tones. It should be noted that switching to a sufficiently high sound level combined with the use of PKM-1 buttons led to a surprising reduction in reaction time, even for an elderly subject, which is why the waiting time range was 1-2000 ms. The results are shown in Table 7.

**Table 7.**
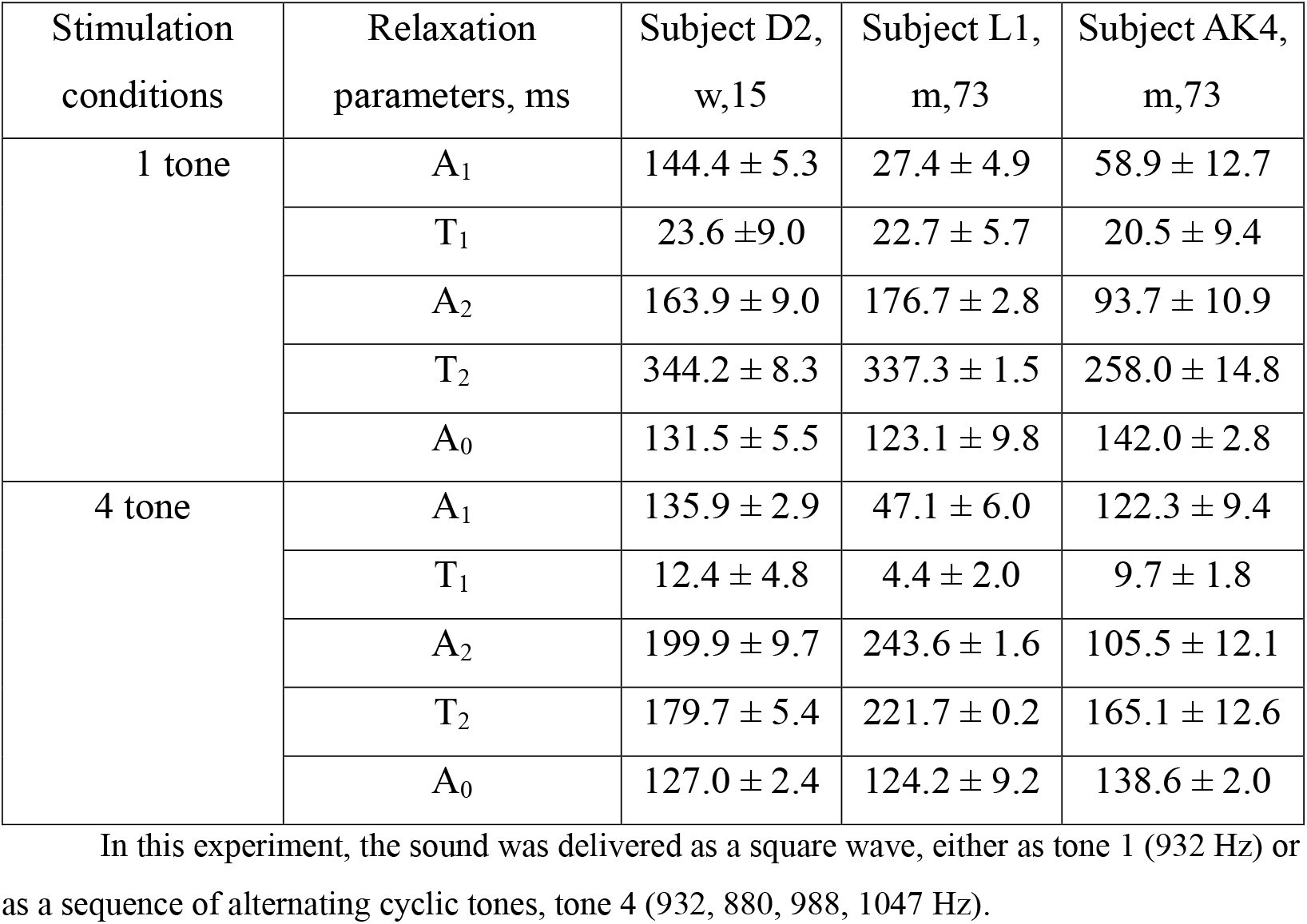
Parameters of a simple audio motor reaction: exposure to single-tone or multi-tone alternating excitation.

The presented data show that the slow relaxation component with alternating stimulation by sounds of different tones accelerates significantly compared to stimulation by a single tone sound.

Moreover, when we tried to change the difference between different sounds, we obtained intermediate values between the 1-sound variant and the “BACH” sounds. This means that the subject distinguished sounds even less than a quarter tone (10 Hz), and a difference of almost a quarter tone (20 Hz) already led to a significant difference from single-tone stimulation.

In the next table, we made a finer comparison, where the distance between cyclically repeating sounds was less than 1 tone. The results are shown in Table 8.

**Table 8.** Dependence of the simple sound-motor reaction (PSMR) parameters on the frequency difference between repeating sounds.

| Relaxation parameters, ms | Difference between tones, Hz |  |  |  |
| --- | --- | --- | --- | --- |
|  | 0 | 10 | 20 | 48-59 |
| A<sub>1</sub> | 98.8 | 122.1 | 184.4 | 114.1 |
| T<sub>1</sub> | 1.7 | 8.3 | 3.85 | 0.8 |
| A<sub>2</sub> | 101.4 | 96.6 | 98.8 | 100.8 |
| T<sub>2</sub> | 248.3 | 235.6 | 219.7 | 201.2 |
| A<sub>0</sub> | 146.5 | 149.1 | 151.5 | 143.7 |

In this experiment, the procedure was the same as for Table 8, except that one base tone ("0") was equal to 932 Hz, and in the rest, the difference between the tones was as indicated, with the exception of the last column, which featured the "BACH" tone set. Subject AK4 (74), (m)

**Table 9.** The effect of alternative stimulation for tactile-motor.

| Relaxation parameters, ms | Stimulating 1 finger | Stimulating 2 fingers alternately |
| --- | --- | --- |
| A<sub>1</sub> | 110,8 ± 8,4 | 125,5 ± 14,7 |
| T<sub>1</sub> | 139,9 ± 28,9 | 38,3 ± 7,4 |
| A<sub>2</sub> | 39,7 ± 6,1 | 78,7 ± 8,2 |
| T<sub>2</sub> | 1479,3 ± 214,1 | 820,3 ± 73,3 |
| A<sub>0</sub> | 147,5 ± 4,5 | 162,5 ± 3,6 |

The excitation was carried out by micro-vibrators (50 ms duration) mounted on the pad of the index finger or alternately on the pads of the index and ring fingers of the subject AK4 (74), (m)

Based on the presented data, it would be interesting to include experiments with a simple tactile-motor reaction. This phenomenon receives much less attention. At the same time, work in this direction is also of interest [32-35]. To elicit the reaction, we used vibrators from smartphones, as they were small in size.

We tested this experiment (once) with 2 more subjects, the result was similar. In this experiment, paired organs were not involved; the fingers were on one hand, but the result did not differ from other variants.

Thus, it can be considered established that stimulation of the simple sensorimotor reaction using alternate excitation of receptors located in different places, or different types of receptors (blue, red color, different sound tones) leads to an acceleration of SSMR relaxation, i.e., the transition to a state of full readiness to respond.

## Discussion

First of all, it should be said that the technique we found – alternating excitation of sensors – has proven its effectiveness. The given imperative, responding to a signal, works independently for different sensors. That is, when one sensor is excited, the other “rests,” restores its ability to send a signal for response to the motor cortex. More precisely, the entire reflex arc from the sensor to the distributor sending the signal to the motor cortex for response “rests.” Moreover, even closely located sensors of different spectral sensitivity for cones, or hair cells, spaced from neighboring ones by 50 Hz, possess this property. This means that when a reflex arc from sensors to the center giving a signal to the motor area of the cortex is created by a verbal imperative, many reflex arcs ending at this center are formed. Upon reaching a certain minimum level, the center issues a signal to the motor area for a motor act (in our case, pressing a button). Upon subsequent excitation, these arcs must first restore their ability to conduct excitation. However, neighboring arcs receiving signals from other receptors can quite satisfactorily send signals to the same center, which gives a signal to the motor area of the cortex. Thus, an impression of apparent acceleration of SSMR relaxation is created.

For light stimulation, we tried to create conditions where the light source fell on the same place on the subject’s retina. This was achieved by a) using one eye and fixing the gaze on a point located approximately 4.1 degrees from the signal LED. The LED size is 5m< m, i.e., about 1 angular degree. Of course, this is not a guarantee that the gaze remains motionless. Nevertheless, light falls both on cones of the required sensitivity and on insensitive cones, which at this moment of the cycle are, as it were, in the dark (referring to cones sensitive to red or blue color). Alternating illumination with red and blue light leads to an apparent twofold decrease in the relaxation time constant of the variable part of the latent time.

This property is characteristic both for sensors with different sensitivity to signal frequency (sound or color) and for sensors located in different areas, albeit quite close, of the retina, i.e., both spatial isolation and frequency isolation of sensors make reflex arcs independent of each other in terms of recovery after excitation.

This observation contradicts the observed effects of diffuse spread of excitation throughout the cortex caused by some event (excitation). Alternatively, it can be assumed that such diffuse spread of excitation occurs after the end of the reflex action or has some other significance for brain function. If it becomes possible to identify various parameters of SSMR relaxation, this approach will allow a more detailed consideration of the reflex arc’s work without external penetration into the brain.

We dealt with a small sample of subjects. Therefore, we cannot claim that all people possess this ability, but at least some people do.

## Notes

### Competing Interest Statement

The authors have declared no competing interest.

## References

1. D.H. Foster Chromatic Function of the Cone: In Encyclopedia of the Eye, Ed D.A. Dartt, AP,// 2010, Pages 266–274 10.1016/B978-0-12-374203-2.00232-3

2. Alekseenko S. V. The structure of the human retina: classical and modern data. Struktura setchatki glaza cheloveka: klassicheskie i sovremennye. // Sensornye sistemy [Sensory systems]. 2019. V. 33(4). P. 269–286 (in Russian). doi: 10.1134/S0235009219040024

3. D. J. Lim Functional structure of the organ of Corti: a review // Hearing Research Volume 22. Issues 1–3. 1986. Pages 117–146 10.1016/0378-5955(86)90089-4

4. D. Deflorio, M. Di Luca and A M. Wing Skin and Mechanoreceptor Contribution to Tactile Input for Perception: A Review of Simulation Models // Front. Hum. Neurosci., 02 June 2022 Sec. Sensory Neuroscience Volume 16, Art..862344 10.3389/fnhum.2022.862344

5. de Haan, E. H., and Dijkerman, H. C. Somatosensation in the brain: A theoretical re-evaluation and a new model. // Trends Cogn. Sci, 2020. 24, 529–541. doi: 10.1016/j.tics.2020.04.003

6. M.M. Murray. J.J. Foxe. B.A. Higgins. D.C. Javitt. C. E. Schroeder Visuo-spatial neural response interactions in early cortical processing during a simple reaction time task: a high-density electrical mapping study// Neuropsychologia 39 (2001) 828–844 10.1016/S0028-3932(01)00004-5

7. C. Escera. E. Yago and K. Alho Electrical responses reveal the temporal dynamics ofbrain events during involuntary attention switching // European Journal of Neuroscience. Vol. 14. pp. 877–883. 2001 DOI: 10.1046/j.0953-816x.2001.01707.x

8. V. Mueller. Y. Brehmer. T. von Oertzen. S.-C. Li and U. Lindenberger Electrophysiological correlates of selective attention: A lifespan Comparison // BMC Neuroscience 2008. 9:18 doi:10.1186/1471-2202-9-18

9. R. DellʼAcqua. P. Dux. B. Wyble. M. Doro. P. Sessa. F. Meconi. and P. Jolicoeur The Attentional Blink Impairs Detection and Delays Encoding of Visual Information: Evidence from Human Electrophysiology// Journal of Cognitive Neuroscience 2015. 27:4. pp. 720–735 doi:10.1162/jocn_a_00752

10. M. J. Ribeiro. J.S. Paiva and M. Castelo-Branco Spontaneous Fluctuations in Sensory Processing Predict Within-Subject Reaction Time Variability// Frontiers in Human Neuroscience 1 May 2016 | Volume 10 | Article 200 doi: 10.3389/fnhum.2016.00200

11. J. Song. C. Cao. Y. Wang. S. Yao. M. P. Catalino. D. Yan. G. Xu and L. Ma Response Activation and Inhibition in Patients With Prolactinomas: An Electrophysiological Study// Frontiers in Human Neuroscience 1 July 2020. Volume 14. Article 170 doi: 10.3389/fnhum.2020.00170

12. A Tlaie. K. Shapcott. T. L. van der Plas. J. Rowland. R. Lees. J. Keeling.. A. Packer. P. Tiesinga M L. Scholvinck. M. N. Havenith What does the mean mean? A simple test for neuroscience// PLoS Comput Biol 2024. 20(4): e1012000. 10.1371/journal.pcbi.1012000

13. S. Gherman. N. Markowitz. G. Tostaeva. E. Espinal. A. D. Mehta. R.G. O’Connell. S.P. Kelly & S. Bickel Intracranial electroencephalography reveals effector-independent evidence accumulation dynamics in multiple human brain regions// Nature Human Behaviour | Volume 8 | April 2024 | 758–770 10.1038/s41562-024-01824-9

14. I.R. Olson. H. Rao. K.S. Moore. J. Wang. J.A. Detre. G. K. Aguirre Using perfusion fMRI to measure continuous changes in neural activity with learning // Brain and Cognition 60 (2006) 262–271 doi:10.1016/j.bandc.2005.11.010

15. O. Baumann. T. Endestad. S. Magnussen. M. W. Greenlee Delayed discrimination of spatial frequency for gratings of diVerent orientation: behavioral and fMRI evidence for low-level perceptual memory stores in early visual cortex // Exp Brain Res (2008) 188:363–369 DOI 10.1007/s00221-008-1366-0J.

16. A. Lewis-Peacock. A.T. Drysdale. K. Oberauer. and B.R. Postle Neural Evidence for a Distinction Between Short-Term Memory and the Focus of Attention // J Cogn Neurosci. 2012 January ; 24(1): 61–79. doi:10.1162/jocn_a_00140.

17. H. Takeuchi. M. Sugiura. Y. Sassa. A. Sekiguchi. Y. Yomogida. Y. Taki. R. Kawashima Neural Correlates of the Difference between Working Memory Speed and Simple Sensorimotor Speed: An fMRI Study // PLoS ONE 2012. 7(1): e30579. doi:10.1371/journal.pone.0030579

18. S. Atmaca. W. Stadler. A. Keitel. D.V.M. Ott. J. ran Lepsien & W. Prinz Prediction processes during multiple object tracking (MOT): involvement of dorsal and ventral premotor cortices // Brain and Behavior 2013; 3(6): 683–700 doi: 10.1002/brb3.180

19. S. E. Paraskevopoulou . W.G. Coon . P. Brunner . K. J. Miller . G. Schalk Within-subject reaction time variability: Role of cortical networks and underlying neurophysiological mechanisms // NeuroImage August 2021. Volume 237. 15. 118127 10.1016/j.neuroimage.2021.118127

20. P.A.F. Laing. T. Steward. C. G. Davey. K.L. Felmingham. M. A. Fullana. B. Vervliet. M. D. Greaves. B. Moffat. R.K. Glarin. and B.J. Harrison Cortico-Striatal Activity Characterizes Human Safety Learning via Pavlovian Conditioned Inhibition// Journal of Neuroscience 22 June 2022. 42 (25) 5047–5057 10.1523/JNEUROSCI.2181-21.2022

21. S.J.S. Isherwood . PL. Bazin . S. Miletić . N.R. Stevenson . A.C. Trutti . D.H.Y. Tse A. Heathcote . D. Matzke . R.J. Innes . S. Habli . D.R. Sokolowski . A. Alkemade . A.K. Håberg . B.U. Forstmann Investigating Intra-Individual Networks of Response Inhibition and Interference Resolution using 7T MRI // NeuroImage 271 (2023) 119988 10.1016/j.neuroimage.2023.119988 .

22. V. Centanino, G. Fortunato and D. Bueti The neural link between stimulus duration and spatial location in the human visual hierarchy// Nature Communications, 2024, 1 5:10720 1 10.1038/s41467-024-54336-5

23. V. Centanino. G Fortunato and D Bueti Defining a functional hierarchy of millisecond time: from visual stimulus processing to duration perception // bioRxiv preprint June 8. 2025 doi: 10.1101/2025.06.06.658257

24. Niemi P., Naatanen R. Foreperiod and simple reaction time // Psychological bulletin. 1981. V. 89. 𝒩 o 1. p. 133. doi 10.1037/0033-2909.89.1.133

25. Los SA, Kruijne W, Meeter M. Outlines of a multiple trace theory oftemporal preparation.// Front Psychol. 2014 Sep 19; 5:1058. (doi:10.3389/fpsyg.2014.01058. eCollection 2014.

26. A.A. Kulakov Features of a Simple Psychophysiological Reaction// Human Physiology, 2018, 44(4):412–417 DOI: 10.1134/S0362119718040060

27. H. Chatroudi, G.Y. Yotsumoto On the nonlinearity of the foreperiod effect // Sci Rep., 2024, Feb 2;14:2780. doi: 10.1038/s41598-024-53347-y

28. A.A. Kulakov Is a Simple Sensorimotor Reaction Really Simple?// BioRxiv preprint may 2 2020 Doi: 10.1101/2020.04.30.070706

29. Kulakov A.A. On the Variability of a Simple Sensorimotor Reaction// Human Physiology, 2023 49(4):364–372 DOI: 10.1134/S0362119722600618

30. A.M. Kukinov,, The use of ordinal statistics and rank criteria for processing observations, // in Poisk zavisimosti i otsenka pogreshnosti (Search for Dependence and Error Evaluation), Moscow: Nauka, 1985, p. 97–110.

31. Kulakov-On the relationship between afterimages and a simple sensorimotor reaction // BioRxiv preprint January 2024 DOI: 10.1101/2024.01.23.576541

32. A. Tomassini, M. Gori, D. Burr, G. Sandini, M. C. Morrone Perceived duration of visual and tactile stimuli depends on perceived speed // Front. Integr. Neurosci., 12 September 2011 Volume 5 - 2011 | 10.3389/fnint.2011.00051

33. S. Reschechtko and J. A. Pruszynski Voluntary modification of rapid tactile-motor responses during reaching differs from its visuomotor counterpart // Journal of Neurophysiology Volume 124, Issue 1 10.1152/jn.00232.2020

34. Sobolev V. V., Popov M. N. Independence of the latency period of a simple tactile-motor reaction from the duration of the tactile stimulus//

35. Scientific Notes jf V.I. Vernadsky Crimean Federal University. Biology. Chemistry, 2024, Volume 10 (76), 𝒩o4, p.196–204 DOI: 10.29039/2413-1725-2025-11-1-170-178

36. V. Sobolev, R. Novak Independence of the latency period of a simple tactile-motor reaction from the preconscious component of sensation during backward masking // Scientific Notes of V I Vernadsky Crimean Federal University Biology Chemistry. 2025, 11(1):170–178 DOI: 10.29039/2413-1725-2025-11-1-170-178

